# *Gynura procumbens* Leaf Extract-Loaded Self-Microemulsifying Drug Delivery System Offers Enhanced Protective Effects in the Hepatorenal Organs of the Experimental Rats

**DOI:** 10.1101/2024.05.15.594312

**Authors:** Manik Chandra Shill, Md. Faisal Bin Jalal, Madhabi Lata Shuma, Patricia Prova Mollick, Md. Abdul Muhit, Shimul Halder

## Abstract

*Gynura procumbens*, known as longevity spinach, is a plant traditionally used in tropical Asian countries for its anti-inflammatory, hepatoprotective, anti-hypertensive, anti-hyperglycemic, and anti-inflammatory properties. The current study aimed to enhance the hepatorenal protective activity of *Gynura procumbens* leaf extract (GLE) by developing a self-microemulsifying drug delivery system (SMEDDS). SMEDDS-GLE exhibited the formation of small micelles with a mean droplet size of 231 nm. This resulted in a significant enhancement in the dispersion of GLE in water, as evidenced by a dispersibility that was at least 4.8 times greater than that of GLE alone. In the rat model of hepatic injury induced by cisplatin (7.5 mg/kg, *i.p.*), the administration of SMEDDS-GLE (75 mg-GLE/kg, *p.o.*) significantly reduced liver damage, observed by histological examination and reduced levels of plasma biomarkers associated with hepatic injury. Furthermore, according to histological examination findings and plasma biomarkers assessment, SMEDDS-GLE enhanced nephroprotective benefits of GLE in the rat model of acute kidney injury. Based on these findings, a strategic application of the SMEDDS-based approach could be a viable choice to enhance GLE’s nutraceutical properties.

## Introduction

In recent years, there has been a significant rise in the diagnosis of kidney dysfunction, including chronic kidney diseases (CKD), among patients with hepatic illnesses ^1^. The growing occurrence of risk factors such as hypertension, diabetes, and nonalcoholic fatty liver disease appears to have had a substantial influence on the increasing occurrence of chronic kidney disease (CKD) ^2^. In 2019, the prevalence of chronic kidney disease (CKD) in hospitalized patients with cirrhosis, the final stage of liver disease, has risen to as high as 46.8% ^3^. Other metabolic risk factors such as obesity, hypertension, and prolonged use of synthetic medicines may be coming together to form a notable epidemiological trend, which may exert the growing prevalence of chronic kidney disease (CKD) among patients with cirrhosis ^4^. Reactive oxygen species (ROS) are significant contributors to pathological alterations in the liver and kidney. Polyunsaturated fatty acids of biological membranes are highly susceptible to the impact of reactive oxygen species (ROS) for lipid peroxidations. Various innate defensive mechanisms have been evolved to restrict the presence of (ROS) and mitigate their detrimental effects ^5,6^. Nevertheless, due to the possibility of incomplete protection or excessive generation of reactive oxygen species (ROS), the inclusion of dietary antioxidants as supplementary protective mechanisms becomes highly significant ^7^. As a result, numerous natural and synthetic substances with antioxidant characteristics have been suggested as potential treatments for liver diseases caused by oxidative stress.

The growing belief that medications derived from natural sources, especially those obtained from food, are more reliable and secure has led to a substantial rise in demand for such goods in the nutraceuticals industry ^8,9^. Functional foods exert therapeutic effects by selectively suppressing the overexpression of proteins, enzymes, amino acids, and hormones through several pathways ^10^. Herbal remedies are gaining popularity in developing and developed nations since they are affordable and have little adverse effects. The World Health Organization (WHO) prioritizes the assessment of secondary metabolites obtained from natural sources for treating diseases, as they have a wide range of effectiveness ^11^. Approximately 80% of the population in the lower income countries relies on natural products to meet their main healthcare requirements. Medicinal herbs are rich sources of exogenous antioxidants, which could be a promising alternative for mitigating pathogenic alterations in oxidative stress-related diseases ^12^. They have been linked to the treatment of liver damage, diabetes, kidney toxicity, cancer, cardiovascular problems, neurological disorders, inflammation, and the aging process. However, the pharmacological potential of these substances may be impeded by their low water solubility, poor dispersibility in the gastrointestinal tract, low stability, poor absorption, and quick metabolism, resulting in a low therapeutic index ^13,14^. Thus, it is inherently challenging to integrate and manage these nutraceuticals within the body for optimum outcomes.

Longevity spinach, scientifically known as *Gynura procumbens*, is a widely recognized traditional food and herb belonging to the Asteraceae family and is commonly used and highly valued in Southeast Asia. *Gynura procumbens* is a plant whose fresh leaves can be consumed raw or cooked ^15^. Multiple reports have demonstrated that *Gynura procumbens* Leaves extract (GLE) contains a variety of secondary metabolites, including quercetin, gallic acid, kaempferol, kaempferol-3-O-β-D-glucopyranoside, kaempferol-3-O-rutinoside, rutin, chlorogenic acid, protocatechuic acid, vanillic acid, syringic acid, caffeic acid, p-coumaric acid, ferulic acid, 3,5-dicafeoylquinate methyl ester, terpenoids, tannins, alkaloids, saponins, and astragali ^16,17^. In addition, it possesses a diverse array of pharmacological actions, including antihyperglycemic, antihyperlipidemic, antioxidant, organ-protective, antibacterial, and anti-inflammatory activity. It is also used for the treatment of hepatotoxicity and alcohol-induced liver disease^18,19^. Many active components (terpenoids, steroids) found in herbal extracts cannot pass through the lipid barrier because they have a high molecular size and low solubility in water ^20^. As a result, they are poorly absorbed into the body and have limited bioavailability.

Nanotechnology has lately emerged as an innovative method to tackle the challenges related to the solubility and bioavailability of medications with low water solubility^16,21^. Several innovative drug delivery methods, such as polymeric nanoparticles, liposomes, nanoemulsions, and phytosomes, have been documented for plant extracts and their bioactive components ^22,23^. Self-microemulsifying formulation improves the absorption of phytomolecules with low solubility ^24,25^. This is achieved by exploiting their lipidic nature and small particle size. A self-microemulsifying drug delivery system (SMEDDS) combines water-insoluble phytoextract, oil or lipid, surfactant, and cosurfactant. Upon being taken orally, they are mixed with water-based fluids in the gastrointestinal tract (GIT) and create a microemulsion/nanoemulsion with droplets ranging in size from 100 to 500 nanometers ^26^. The microemulsion is designed to dissolve the phytomolecules necessary to absorb, which are not easily soluble in water. In addition, the lipid excipients in the self-microemulsifying formulation enhance the transportation of phytomolecules through the lymphatic system, improving bioavailability by reducing the effects of first-pass metabolism. The intracellular concentration of phytomolecules increases due to the weakening of the lipid and surfactant employed by the *P*-glycoprotein efflux mechanism ^27^. By boosting the biopharmaceutical properties and enhancing the stability of the gastrointestinal system, the employment of the SMEDDS-based method may help overcome the challenges associated with giving GLE in clinical settings. Therefore, the objective of the present study is to create, analyze, and assess SMEDDS for the oral administration of GLE. The purpose is to enhance the biopharmaceutical properties of GLE, leading to improved hepatoprotective and nephroprotective effects.

## 1. Materials and Methods

### 1.1. Chemicals and Reagents

BASF, Dhaka, Bangladesh graciously gave Kolliphor^®^ P188 and medium chain triglyceride (MCT). Cisplatin was bought from Sigma-Aldrich (St. Louis, MO, USA). Throughout the experiment, all other reagents and solvents were of analytical grade and were acquired from commercial sources.

### 1.2. Collection, Identification, and Extraction of Plant Materials

The National Herbarium in Mirpur, Bangladesh, identified and verified *Gynura procumbens* leaves collected from Natore, Bangladesh. A voucher specimen (DACB accession number: 45273) was recorded at the herbarium. After gathering the leaves, they were carefully cleaned, dried in the shade, and powdered into a fine powder. The leaf powder was soaked in ethanol, thoroughly agitated, filtered, and dried to a crude extract using a rotary evaporator as described in our previous study ^28^. To use later, the extract was kept in an airtight falcon tube and refrigerated at 2 to 8 °C.

### 1.3. HPLC detection and quantification of polyphenolic compounds

The study of specific phenolic components in the ethanol leaf extract of *Gynura procumbens* was carried out using HPLC-DAD detection and quantification techniques. A Dionex UltiMate 3000 system with a photodiode array detector (DAD-3000RS) and quaternary rapid separation pump (LPG-3400RS) was used. An Acclaim^®^ C18 (5 µm) Dionex column (4.6 × 250 mm) at 30 °C, 1 mL/min flow rate using gradient elution mode, and 20 µL injection volume were used to achieve separation. Acetic acid solution pH 3.0 (solvent B), methanol (solvent C), and acetonitrile (solvent A) made up the mobile phase. The detector was calibrated to measure 280 nm for 25.0 min, then 320 nm for 32.0 min, 280 nm for 3 min, and 380 nm for 35 min. The diode array detector was set to an acquisition range of 200 to 700 nm and maintained for the remainder of the analysis. A standard stock solution was made in methanol to create calibration curves. It contained the following: kaempferol (6 µg/mL), pyrogallol, rutin hydrate (12 µg/mL), p-coumaric acid (3 µg/mL), ellagic acid (48 µg/mL each), vanillin, trans-ferulic acid (+)-catechin hydrate, (-)-epicatechin (10 µg/mL each), and quercetin hydrate (QU) (4 µg/mL). The extract was dissolved in ethanol at a 30 mg/mL concentration. Before HPLC analysis, all solutions mixed standard, sample, and spiked solutions were degassed in an ultrasonic bath (Soner 210H, Rocker Scientific, Taiwan) for 15 min after being filtered through a 0.22 µm syringe filter (Sartorius, Germany). Dionex Chromeleon software was used to calculate data acquisition, peak integration, and calibrations (Version 6.80 RS 10) ^28^.

### 1.4. Formulation of SMEDDS-GLE

By combining various ratios of oil, surfactant, and co-surfactant with the amount of GLE (5%, w/v), a series of SMEDDS-GLE were made. In summary, oil, surfactant, and co-surfactant were accurately measured in glass vials, mixed by inverting, and then heated in a water bath at 37 °C while stirring moderately. The ultimate blend was agitated using a vortex mixer until a transparent solution was achieved. Before being used, the resulting mixture was kept at room temperature in an amber glass bottle.

### 1.5. Preparation of pseudo-ternary phase diagram

Using the water titration method, the pseudo-ternary phase diagrams of oil, surfactant, and co-surfactant were created. The oil, surfactant, and co-surfactant mixtures at different weight ratios were combined with water drop-wise, and the resulting emulsion formation was assessed immediately. The mixtures were then gently shaken with water, and the dispersibility, phase clarity, flowability, and precipitation rate were visually evaluated. Following phase diagram validation of the fine emulsion region, the appropriate oil, surfactant, and co-surfactant ratio was chosen for further physicochemical and pharmacodynamic analyses. Based on the observed data, the pseudo-ternary phase diagram was produced using SigmaPlot V14 (Systat Software Inc., UK).

### 1.6. Interaction of GLE with Polymers

The compatibility and potential interactions between the components of SMEDDS-GLE were assessed by Attenuated Total Reflectance-Fourier Transform Infrared (ATR-FTIR) analysis using a Shimadzu IR-Prestige-21-FT-IR infrared spectrometer (Tokyo, Japan) connected to a horizontal Golden Gate MKII single-reflection ATR system (Specac, Kent, UK). Each sample was placed separately on the instrument’s sample platform. For each spectrum, 64 consecutive scans were collected in the 600–4000 cm^−1^ (with a resolution of 4 cm^−1^). The final spectrum was generated by averaging these observations. The IRsolution 1.30 software (Shimadzu, Tokyo, Japan) was used for all spectral analyses. During the analysis, the baseline of each sample was normalised and corrected.

### 1.7. Dynamic Light Scattering (DLS)

Using a Malvern Zetasizer Ultra (Malvern Instruments Ltd, Malvern, UK), the DLS method was used to assess the mean particle size, polydispersity index (PDI), and zeta potential of SMEDDS-GLE samples suspended in water at 25 °C and a measuring angle of 90°. After being ten-fold diluted with Milli-Q water, the samples were used to calculate the zeta potential and size distribution. Three replicates of each measurement were carried out.

### 1.8. Transmission Electron Microscopy (TEM)

Using TEM, the droplets of the optimized SMEDDS-GLE formulation were investigated. In particular, SMEDDS-GLE was poured onto a 300-mesh copper grid coated with carbon after being diluted 400 times with Milli-Q water (0.5%, w/v). Using Milli-Q water, excess phosphotungstic acid (PTA) was eliminated following negative staining with a 2% (w/v) PTA solution. After that, the copper grid was put in an oven to remove all traces of moisture through vacuum drying. At a voltage of 200 kV, the dried grid was examined using TEM equipment (F200X Talos, Thermo Fisher, Waltham, MA, USA). The droplet size was verified using ImageJ (National Institutes of Health, Bethesda, MD, USA).

### 1.9. *In vitro* dissolution/dispersion tests

A study assessed the initial self-emulsifying ability of GLE samples (1 g-GLE) by *in vitro* dissolution/dispersion testing. The study employed the USP Type-II dissolution test apparatus, distilled water, and a pH 1.2 (0.1N HCl) solution as the testing mediums. The dispersion test was conducted for 60 min in 900 mL of the media at 37 ±0.5°C, with continuous stirring at a speed of 50 rpm. Subsequently, the samples were extracted from the central location of the dissolving vessel at specified time intervals (5, 15, 30, 45, and 60 min) using a micropipette. Following each sampling, an equivalent volume of fresh media has been added to the vessel. Subsequently, the sample was centrifugated at 10,000 × *g* for 10 min. The resulting mixture was then passed through a 0.45 µm membrane filter (Millex LG, Millipore, Billerica, MA) and diluted with methanol. The UV spectrophotometer was used to analyze the amounts of dispersed or dissolved GLE at a wavelength of 254 nm ^29^.

### 1.10. Animals

Animal models play a vital role in evaluating fundamental pharmacokinetic and pharmacodynamic factors, such as drug metabolism, effectiveness, safety, and toxicological research, before conducting human trials. Selecting an appropriate animal model can be a difficult task that typically involves making assumptions and considering the feasibility of the study ^30^. Rodents are a vital animal species for biomedical research. Mice and rats comprise more than 95% of the animals used in biomedical research. Their physiology and genetic makeup are almost identical to that of humans, making them a suitable choice for scientific research ^31^. Locally bred healthy Wistar albino rats (male rats, body weight within 250−300 g) were obtained from North South University, Dhaka, Bangladesh, and kept in pairs in cages. They had unrestricted access to food and water and were subjected to a 12 h cycle of darkness and light. The environment was maintained at a controlled temperature of 24 ± 1 °C and humidity of 55 ± 5% RH. Before the oral delivery of GLE samples, the rats underwent a period of food deprivation lasting at least 12 h, excluding water. All animal-related procedures in this study were carried out per the guidelines approved by the Institutional Animal Care and Ethical Committee of the Faculty of Biological Sciences, University of Dhaka (Approval No. 233, 30 August 2023). In addition, our experiments also adhered to the International Council for Laboratory Animal Science, the Nuffield Council on Bioethics (NCB), and the Council for International Organization of Medical Sciences (CIOMS/ICLAS) guidelines.

### 1.11. Experimental rat model of Kidney and Liver Injury

In a rat model of acute kidney and liver injury induced by cisplatin ^32^, the nephroprotective and hepatoprotective properties of GLE samples were assessed. A single intraperitoneal injection of cisplatin (7.5 mg/kg) dissolved in saline was given to rats to cause immediate damage to the kidneys and liver. Rats were randomly divided into four groups, each with six rats: group I was the control group (received saline); Group II was the disease group (received cisplatin); Group III was the GLE treatment group (received cisplatin and GLE); and group IV was the SMEDDS-GLE treatment group (received cisplatin and SMEDDS-GLE). All rats, except those in groups I and II, received treatments (both GLE and SMEDDS-GLE) for 10 days and an intraperitoneal injection (IP) of cisplatin (7.5 mg/kg dissolved in saline) on the 7^th^ day of the experiment and the treatment with GLE samples continued till 10^th^ day. The dose of GLE and SMEDDS-GLE was 75 mg/kg to assess the nephroprotective and hepatoprotective effects based on the findings of exploratory tests based on earlier reports ^28^. At the end of the experiment, rats were anesthetized using ketamine (100 mg/kg), blood samples were collected, centrifuged at 8,000 × g for 10 min, and plasma was refrigerated at -80 °C until analysis. The rats were euthanized using ketamine (300 mg/kg) and sacrificed to collect kidney and liver tissues. The collected tissues were rinsed with ice-cold saline, dried with filter paper, and well-preserved (10% neutral formalin) for further analysis. After two days of cisplatin (6 mg/kg), rats showed increased thirst, weight loss, anemia, decreased activity, and gastrointestinal distress ^33^. Gastrointestinal malaise is characterized by reduced food and drink intake and impaired stomach function, as indicated by a notable increase in gastric content. Following sample collection, a humane endpoint was implemented to prevent the rats from experiencing pain or distress, and they were subsequently euthanized ^34^. After being treated with cisplatin, all the rats lived through the whole study.

### 1.12. Evaluation of plasma biochemical markers

BUN and creatinine levels were assessed as indicators of nephrotoxicity using LiquiUV diagnostic kits, a commercially available test kit (HUMAN GmbH, Germany). Following the manufacturer’s instructions, the creatinine and BUN levels in plasma were measured spectrophotometrically at 520 and 340 nm using a microplate reader (Safire, Tecan, Männedorf, Switzerland). Moreover, the Fortress diagnostic kit (UK) was used to measure the plasma levels of alkaline phosphatase (ALP), aspartate aminotransferase (AST), and alanine aminotransferase (ALT) as markers of liver impairment. The ALT and AST levels were measured spectrophotometrically at a wavelength of 340 nm. Every sample was assessed three times.

### 1.13. Histopathological examination of liver and kidney

The liver and kidney tissues were subjected to histological scanning following the methodology described by Quaresma et al. with slight modifications ^35^. The collected tissues were then preserved in 10% neutral buffered formalin. The fixed tissues were rinsed three times with phosphate-buffered saline (PBS, pH 7.4) and then placed in a solution of 30% sucrose and 0.1% sodium azide at 4 °C for 24 h. The kidney and liver tissues were sliced into sections with a thickness of 5 µm using a rotary microtome and preserved in paraffin wax. Hematoxylin and eosin (H&E) staining was used to investigate the morphology of the liver and kidney tissues and the presence of inflammatory cell infiltration to evaluate the severity of tissue damage. Subsequently, the sections were carefully investigated, recorded, and photographed using a Zeiss Axioscope 40-X light microscope.

### 1.14. Data analysis

The findings are presented as the mean ± standard error of the mean (SEM). The data was subjected to a one-way analysis of variance (ANOVA) using GraphPad Prism-6 software (San Diego, California, USA; http://www.graphpad.com) to confirm its statistical significance. Statistical differences were considered significant at a *p* ≤ 0.05.

## 2. Results and Discussion

### 2.1. Characterization of Polyphenolic Compounds in the Ethanolic Leaf Extract of *Gynura procumbens*

The phenolic phytochemicals in the ethanolic leaf extract of *Gynura procumbens* were detected and measured using HPLC. **Fig. 1** displays the chromatographic separations of polyphenols in *Gynura procumbens*. The findings of the investigation on the phenolic chemical found in GLE are shown in **Table 1**. The quantities were determined using the relevant calibration curve and presented as the three measurements’ averages.

**Fig. 1.**
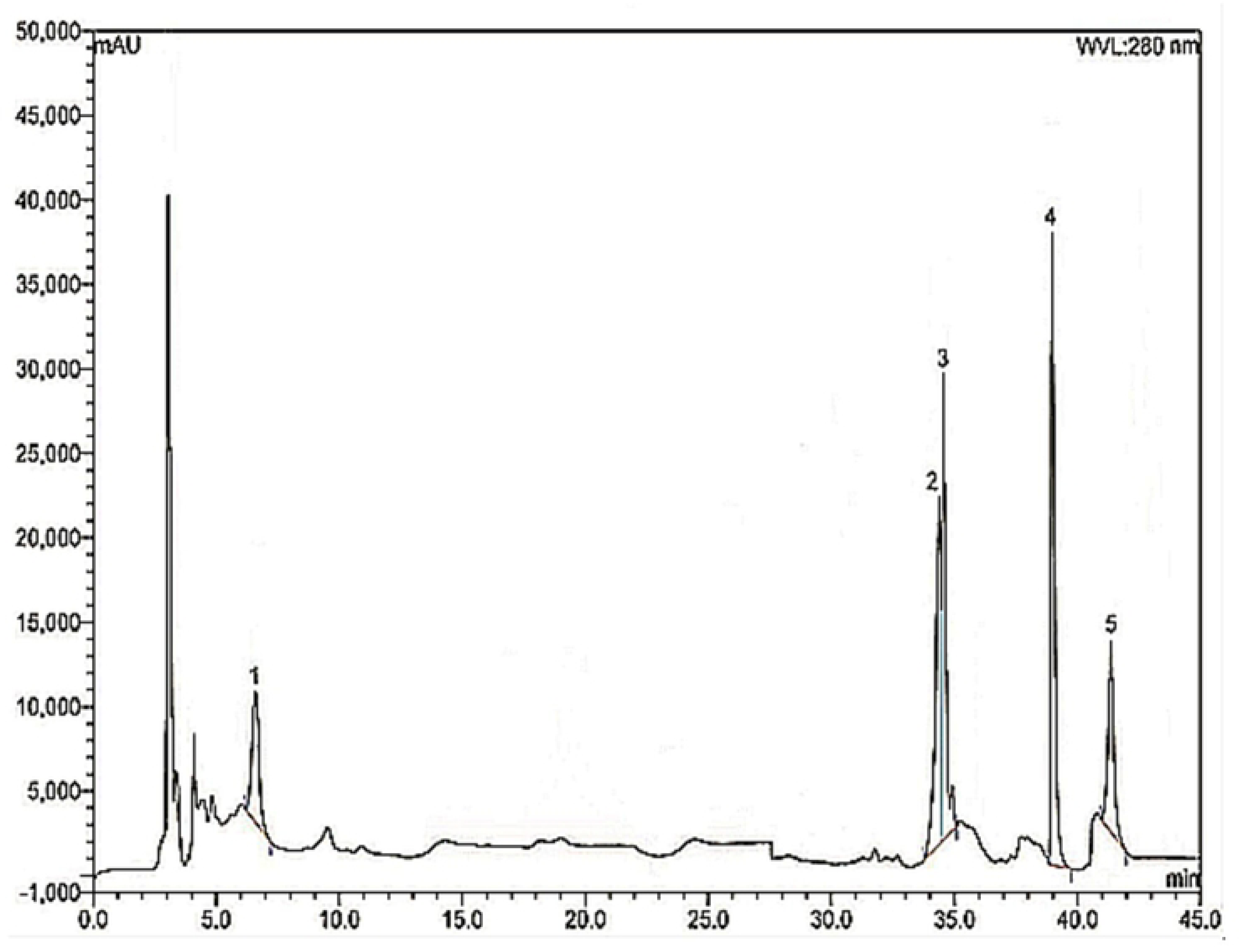
HPLC chromatogram of ethanolic extract of *Gynura procumbens*

**Table 1.** Polyphenolic compounds identified in the ethanol leaf extract of *G. procumbens*.

| Polyphenolic compounds | Ethanollic extract of *Gynura procumbens* |  |
| --- | --- | --- |
|  | Content (mg/100 g of dry extract) | %RSD |
| Gallic acid | 10.24 | 0.38 |
| Ellagic acid | 22.09 | 0.54 |
| Rutin hydrate | 40.52 | 0.71 |
| Quercetin hydrate | 58.27 | 0.92 |
| Kaempferol | 13.48 | 0.47 |

RSD: Relative standard deviation (n = 3)

### 2.2. Optimization of SMEDDS composition

SMEDDS containing oil, surfactant, and co-surfactant are monophasic liquids at room temperature; however, when mixed with water, they form a fine oil-in-water emulsion upon gentle stirring. The solubility of the target molecules in these components is critical for creating a clear, monophasic liquid at room temperature and maintaining it solubilized. Emulsification depends on oil, water, surfactant, and cosurfactant/cosolvent proportions. In the absence of GLE, pseudo-ternary phase diagrams (PTPD) were used to locate self-emulsifying regions and optimize SMEDDS oil, surfactant, and cosurfactant concentrations. The PTPD determines the correct oil, surfactant, and cosurfactant/cosolvent ratio for emulsion with varying water volumes. Different ratios of oil, surfactant, and co-surfactant containing SMEDDS were created, and spontaneous self-emulsification was visually inspected to determine the fine emulsion areas and optimize the ratios (**Fig. 2**). The isotropic mixture of oil, surfactant, and co-surfactant creates an O/W emulsion in water when gently stirred. The change in entropy favors dispersion more than the energy needed to increase its surface area, causing self-emulsification. Oil facilitates medication transport across the intestinal lymphatic system, improving absorption and bioavailability in SMEDDS.

**Fig. 2.**
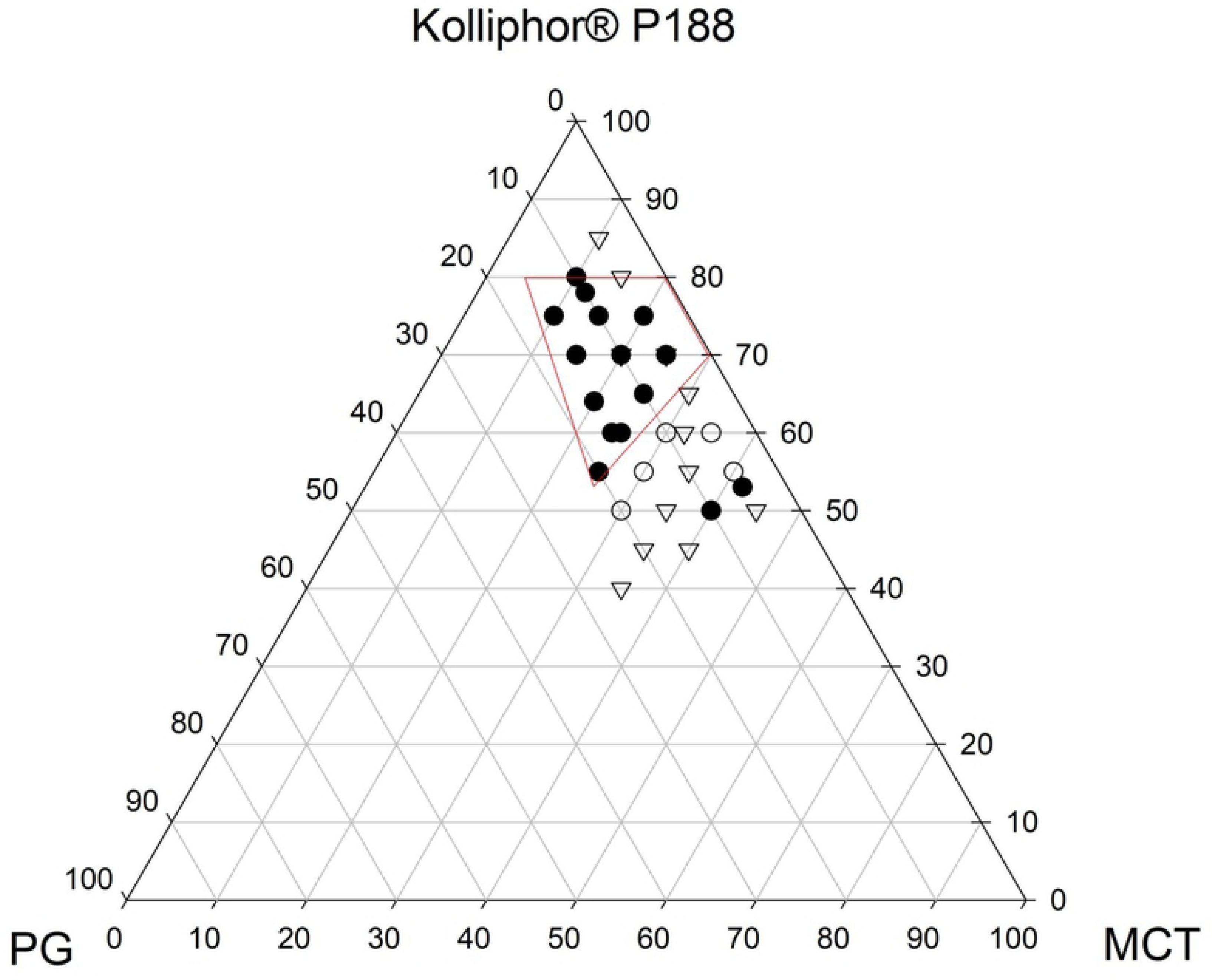
Pseudo-ternary phase diagram of SMEDDS formulations of GLE in water containing surfactant (Kolliphor^®^ P188), oil (MCT), and co-surfactant (PG). ● (filled area), fine emulsion region; ▽, coarse emulsion region; and ◯, poor or not emulsified.

Additionally, oil-phase fatty acids can affect self-emulsifying system features. Short-chain triglyceride (SCT) or medium-chain triglyceride (MCT) oils also help in nanoemulsion formation. For SMEDDS oil phases, a fine drug solubilizing oil is recommended to maximize target compound loading and prevent drug precipitation following dispersion. Additionally, MCT exhibits superior emulsification compared to long-chain glycerides. The study utilised Kolliphor^®^ P188, a non-ionic amphiphilic surfactant licensed for pharmaceutical and food usage, due to its low toxicity (LD_50_ = 9380 mg/kg in a rat study) and ability to stabilise emulsions ^36^. It was found that at least 50% w/w Kolliphor^®^ P188 was needed to disperse lipids and generate fine emulsion ^37^. According to research findings, co-surfactants are crucial components in SMEDDS formulations that boost target compound loading and enlarge the self-emulsification zone in phase diagrams ^38^. A single surfactant rarely provides the fluid interfacial film, and microemulsion stability requires brief negative interfacial tension ^39^. Co-surfactants usually reduce interface bending stress and give the interfacial layer enough elasticity/flexibility to generate microemulsions in various compositions. This investigation used propylene glycol (PG) as a co-surfactant to stabilise emulsions and finely solubilize GLE. PG is an approved food excipient and commonly utilized in nutraceutical formulations due to its low toxicity (LD_50_ = 44 g/kg in rat, oral) ^40^. PG, as a co-surfactant, might enhance dispersibility by lowering oil-water interfacial tension, potentially accelerating drug absorption in the gastrointestinal system. Increasing surfactant concentration improved self-emulsification, but surfactant concentrations below 50% resulted in poor emulsification, as previously observed. Increasing surfactant concentration stabilizes the oil-water interface, forming a thin, low-viscosity emulsion with microscopic droplets. Higher doses of surfactant and co-surfactant stabilize emulsions with lower globule diameters, resulting in finer emulsions. To minimize toxicity and tissue simulation potential, minimal amounts of surfactant and co-surfactant were selected in formulations for *in vivo* use. A 20:65:15 ratio of oil, surfactant, and co-surfactant was chosen for optimal SMEDDS formulation of GLE, as well as for physicochemical and *in vivo* assessments.

### 2.3. Physicochemical characterizations of SMEDDS-GLE

The efficiency of fine self-emulsification is strongly dependent on droplet size upon exposure to aqueous media to form fine submicron dispersion, and the stability of SMEDDS varies on droplet size distribution, according to the liquid formulation classification system, type-III ^41^. DLS analysis (**Fig. 3 A**) and TEM observation (**Fig. 3B**) were used to characterize the emulsification property of SMEDDS-GLE products. DLS data showed fine emulsion droplets with an average particle size of about 231 nm and a polydispersity index (PDI) of 0.12 following the dispersion of SMEDDS-GLE in water. Furthermore, TEM investigations revealed that many spherical emulsions with a particle size of roughly 200 nm made up the dispersed SMEDDS-GLE in water. The self-emulsified GLE exhibited a negative surface charge, as indicated by a zeta potential of approximately -48.9 mV. This negative charge could be attributed to free fatty acids on the surface of emulsions derived from MCT, which may prevent particle aggregations due to the electric repulsion between particles ^42^. As per a prior study, colloidal dispersions exhibit favorable stability when their zeta potential is high and negative, exceeding −20 mV ^43^. This is because of the electric repulsion between the particles. The great colloidal stability of the formulation was also demonstrated when SMEDDS-GLE was dispersed in several aqueous media, such as water, SGF, and SIF, at room temperature, and at least three hours later, no discernible changes in the particle size or zeta potential occurred. There was a small difference between the particle sizes in the DLS study and the TEM examination. The disparity between the two approaches is most likely the cause of this variance. While TEM examination is performed on dry samples where the corona is squeezed on the droplet surface, DLS measures a hydrodynamic size where the hydrated corona of the stabilizing polymer contributes significantly to the observed diameter. These results imply that the SMEDDS-GLE had a comparatively narrow size distribution and good dispersibility, which may have improved GLE’s oral absorption and dissolving characteristics.

**Fig. 3.**
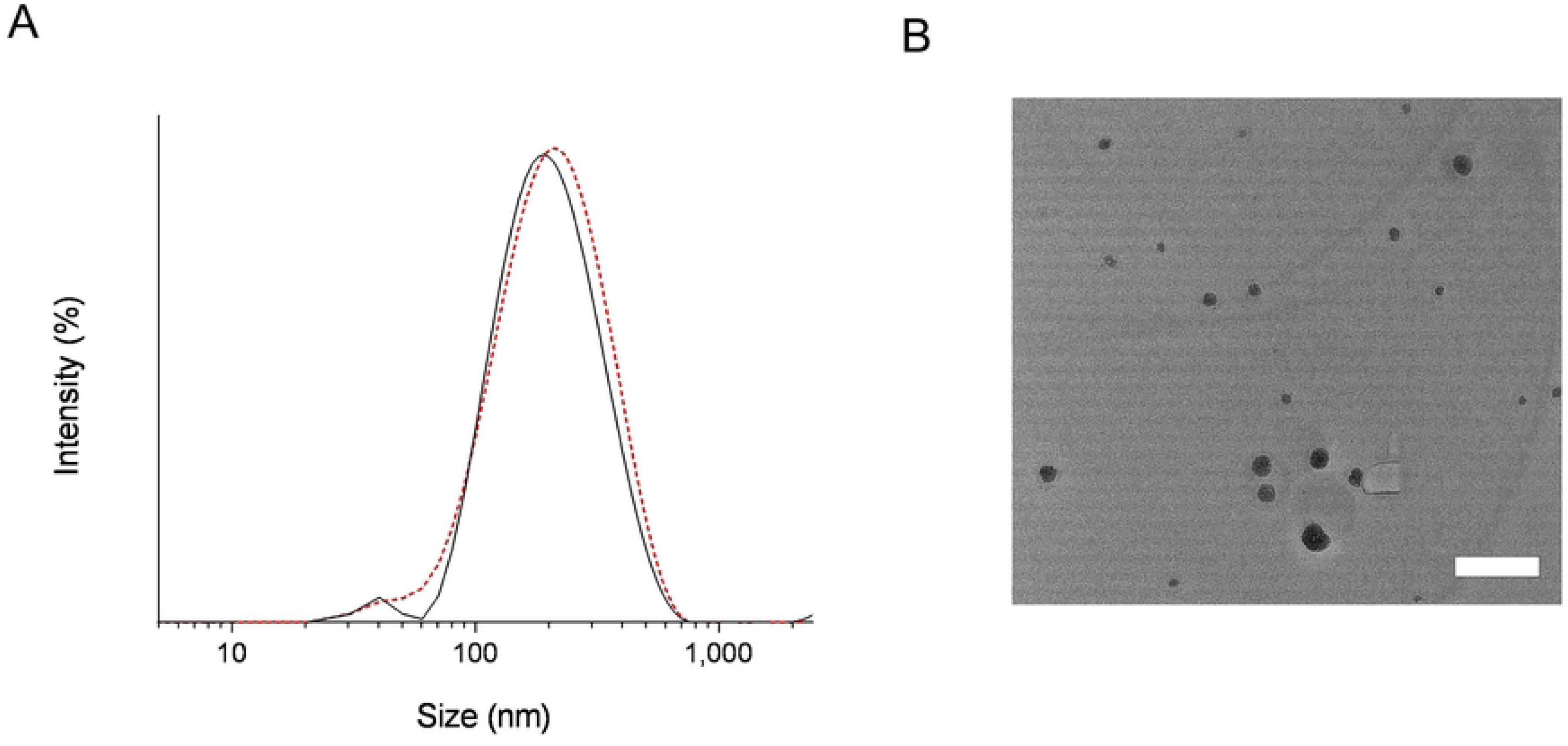
Micelle forming potency of TQM samples dispersed in water. (A) DLS analysis determined the emulsion size distribution of SMEDDS-GLE spread in distilled water: The solid line represents the micelle size distribution at 0 min after dispersion, and the dotted line represents the micelle size distribution at 3 h after dispersion. (B) TEM image of SMEDDS-GLE dispersed in distilled water. Bar represents 200 nm.

### 2.4. *In vitro* dispersion/dissolution test of SMEDDS-GLE

. The dissolution/dispersion behavior of GLE samples was shown in **Fig. 4** Improved clinical outcomes are indicated by increased dissolution/dispersion, which indicates a higher dissolving rate followed by higher absorption. Poor GLE dispersion was indicated by the 18.8% dissolution/dispersion in water after 60 min. On the other hand, at least 4.8-fold better dissolution/dispersion behavior was seen in SMEDDS-GLE, suggesting fine emulsification (**Fig. 4A**). However, in an acidic environment, GLE showed poor dissolution/dispersion behavior, with 13.3% of the GLE dispersed after 60 min. In contrast, SMEDDS-GLE showed better dispersion, increasing by 7-fold after 60 min before plateauing (**Fig. 4B**). Rapid GLE dissolving and dispersion behaviors in SMEDDS-GLE could be demonstrated from the dissolution/dispersion behavior data, and this SMEDDS-GLE technique would accelerate the high and quick oral absorption of GLE’s active ingredients. The dissolution rate of the SMEDDS was significantly enhanced by the reduced particle size and enlarged surface area of the emulsion droplets. Including surface stabilizers in plant, extract enhances the solubility and wettability of poorly soluble flavonoids in the dissolving medium. This, in turn, contributes to the faster dissolution rate of SMEDDS-GLE.

**Fig. 4.**
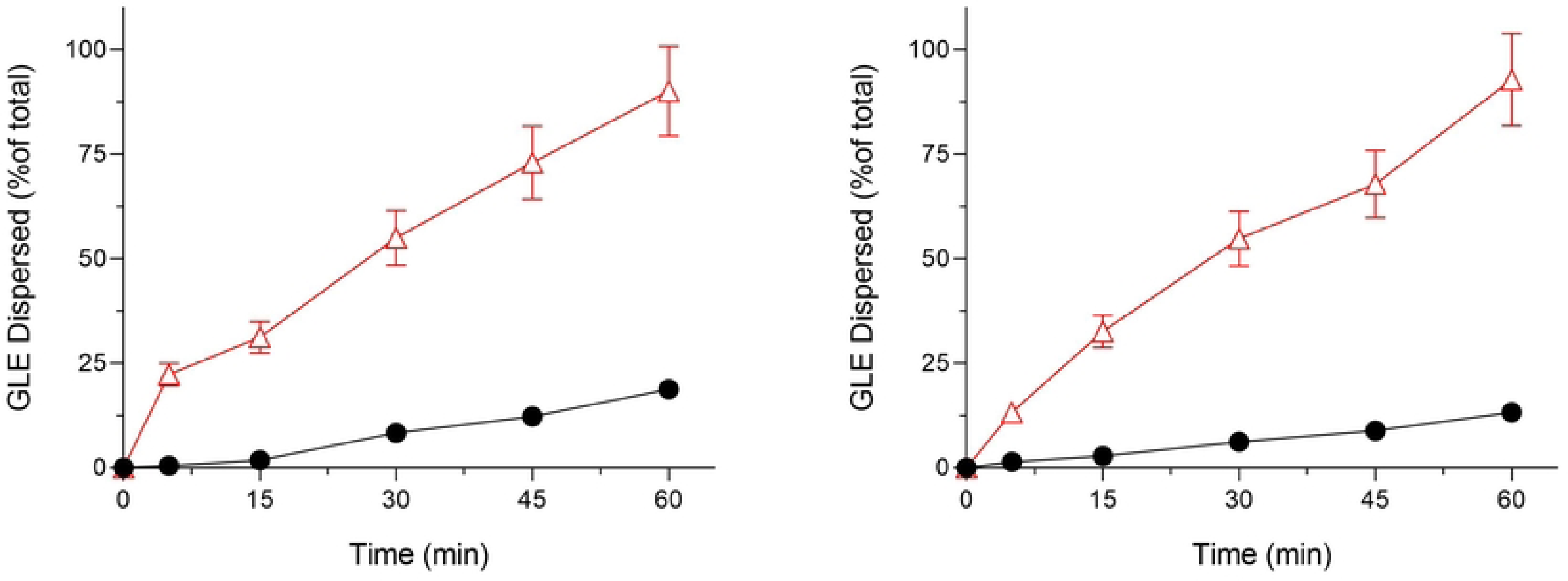
Dissolution/dispersion tests of GLE samples (A) Dissolution/dispersion tests in distilled water; (B) Dissolution/dispersion tests under acidic conditions (0.1 N HCl, pH 1.2). ●, GLE and △, SMEDDS-GLE. Each bar represents the mean ± S.D. of 3 independent experiments.

### 2.5. Interaction among SMEDDS-GLE components using FTIR-ATR analysis

The FTIR-ATR study was conducted to determine the possible interaction between the components of SMEDDS and GLE. GLE is known for its abundance of quercetin, gallic acid, kaempferol, rutin, chlorogenic acid, protocatechuic acid, vanillic acid, syringic acid, caffeic acid, p-coumaric acid, and ferulic acid, which are believed to contribute to its pharmacological effects. The FTIR spectra analysis has identified a variety of distinct peaks that can be attributed to different functional groups, such as C-H, -CH_2_, -CH_3_, C=O, and C=C (**Fig. 5**). The spectra of GLE showed distinct absorption bands at 1370 and 1167 cm^−1^, indicating the presence of C-H bending and C-O stretching groups in the GLE. Furthermore, the presence of methyl and isopropyl substituents is indicated by the two strong bands seen at 2,923 cm^−1^ and 2,853.6 cm^−1^, which can be attributed to the stretching of C-H bonds in an aliphatic group. Additionally, a significant and distinct band is detected at 1,743.7 cm^−1^, indicating the presence of the ketone group through stretching the C=O bond. The two additional peaks observed at 1,463.8 and 1,377.5 cm^−1^ can be attributed to the absorption of C-H bonds in a scissoring motion and the vibration of methyl groups, respectively. However, there was a small change in the bands in the C=O region (1,650–1,750 cm^−1^) in SMEDDS-GLE.

**Fig. 5.**
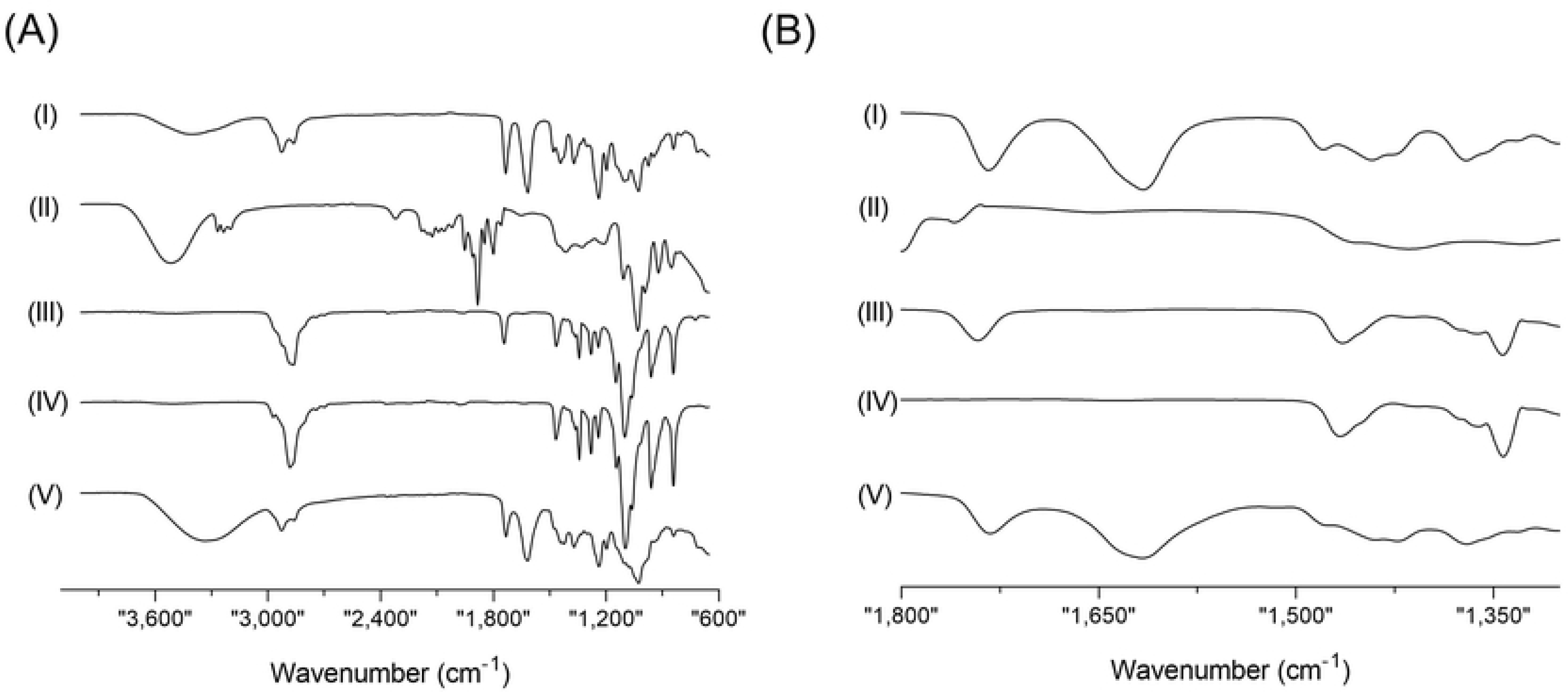
ATR-FTIR spectroscopic analysis of GLE samples in the spectral region from (A) 4,000–600 cm^−1^ and (B) 1,800–1,300 cm^−1^. (I) SMEDDS-GLE, (II) PG, (III) MCT, (IV) Kolliphor^®^ P188, and (V) GLE.

Furthermore, in the SMEDDS-GLE system, the spectra exhibited a faint and widened peak in the C-H stretching region, attributed to the hydrophobic interaction. The results of the FT-IR spectrum analyses suggest a modest conformational transformation in the spectra, indicating that GLE was dispersed in SMEDDS. This dispersion led to enhanced dissolution and dispersion behavior, as observed in previous studies ^44,45^.

### 2.6. Hepatoprotective Effects of SMEDDS-GLE

Multiple published studies have shown that natural antioxidants, such as polyphenols, are powerful bioactive antioxidants that disrupt lipid peroxidation chains by neutralizing highly reactive oxygen radicals. Studies have demonstrated that GLE can enhance the levels of hepatic endogenous antioxidant enzymes such as superoxide dismutase and catalase. This effect may be beneficial in protecting against oxidative diseases caused by free radicals. The study found that SMEDDS-GLE enhanced the biopharmaceutical properties of GLE. As a result, SMEDDS-GLE may demonstrate enhanced hepatoprotective benefits against hepatic injury induced by cisplatin in rats. Plasma ALT, AST, and ALP values as surrogate biomarkers were considered to assess hepatotoxicity and evaluate hepatocellular damage in a rat model treated with cisplatin. The findings demonstrated a significant increase in ALT, AST, and ALP levels in rats treated just with cisplatin compared to the control group. This suggests considerable damage to the liver tissues (**Fig. 6**).

**Fig. 6.**
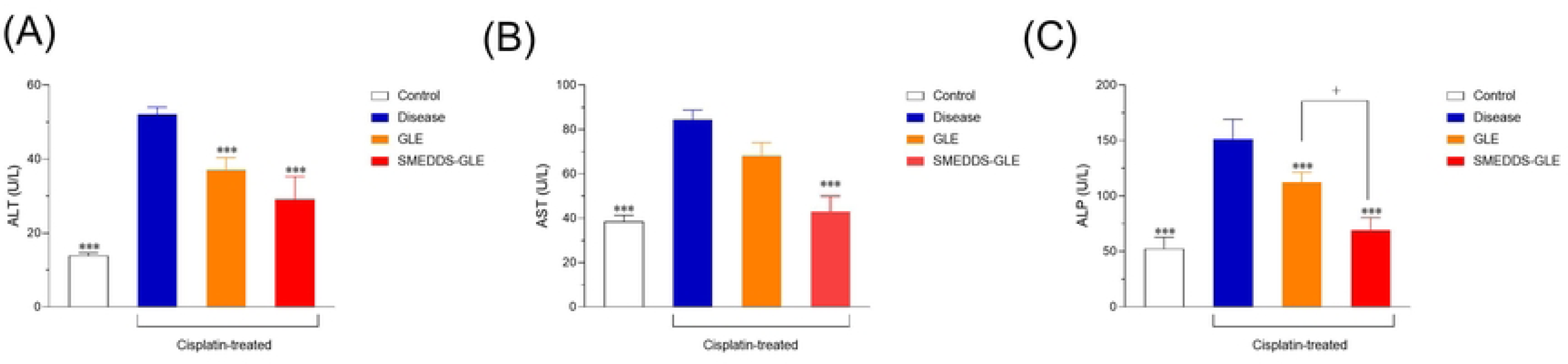
Effect of GLE samples on hepatic markers in the rat liver: (A) Aspartate Amino Transferase (AST), (B) Alanine aminotransferase (ALT) and (C) Alkaline Phosphatase (ALP) levels. The data were expressed as Mean ± S.E.; n=6, where n is the number of determinations. **, *P*<0.01; ***, *P*<0.001 with respect to cisplatin-treated rats; ##, *P*<0.01; ###, *P*<0.001, GLE *vs.* SMEDDS-GLE. Data represent mean ± S.E. of 6 experiments.

Treatment with GLE (75 mg/kg, *p.o.*) in rats reduced ALT levels by 28.9%, AST levels by 19.2%, and ALP levels by 25.8% compared with the cisplatin-treated group alone. On the other hand, treatment with SMEDDS-GLE (75 mg-GLE/kg, *p.o.*) provides a significant reduction in ALT, AST, and ALP levels by 44.2%, 49.05%, and 54.2%, respectively (**Fig. 5**). This can be attributed to the minimal absorption of bioactive components from GLE through oral administration, as shown in **Fig 6**. Hepatic ALT is located in the liver and is vital in catalyzing the transformation of proteins into energy, particularly for hepatocytes ^46^. The level of this substance in the plasma increases when the liver is injured since it is released into the bloodstream. However, it has been shown that AST plays a crucial part in the metabolic processes related to amino acids ^47^. Like ALT, AST is commonly detected in the bloodstream at relatively low concentrations. Increased AST levels may suggest a pathogenic disease, liver damage, or muscle injury ^27,48^. ALP are ectoenzymes linked to plasma membranes that hydrolyze blood monophosphate esters, aiding in the breakdown of proteins ^49^. An ALP test determines the amount of ALP circulating in the blood.

One of the tests in a comprehensive metabolic panel is the measurement of ALP levels, which can be high or low and suggest an underlying disease originating from the liver and bones ^50^. The study revealed elevated ALP, AST, and ALT levels in the groups treated with cisplatin, suggesting potential hepatotoxicity in the animals. On the other hand, SMEDDS-GLE extract effectively decreased the levels of these enzymes which might be beneficial in correcting the liver tissue abnormalities induced by cisplatin in rats. The results indicate that the improved distribution of GLE in the liver using SMEDDS-GLE helped boost the hepatoprotective effects of GLE. Histological assessment frequently utilizes hematoxylin and eosin (H&E) staining since it offers pathologists and researchers a comprehensive visual depiction of the examined tissue. To achieve this purpose, distinct staining techniques are used to visualize and differentiate various cellular components, such as the cytoplasm, nucleus, organelles, extracellular components, and other important characteristics of cells ^51^. The histological investigation of H&E-stained liver tissues, as depicted in **Fig. 7**, revealed that the control group exhibited distinct cellular architectures characterized by robust cytoplasm and healthy hepatic cells. On the other hand, the liver tissues of the group treated with cisplatin exhibited noticeable harm, including substantial fatty alterations (hepatic steatosis), fibrotic areas between the nodules, necrosis, and the deterioration of parenchymal cells (**Fig. 7B**). These findings indicate that SMEDDS-GLE has the potential to reduce liver cell damage and could be advantageous in preventing illnesses caused by free radicals.

**Fig. 7.**
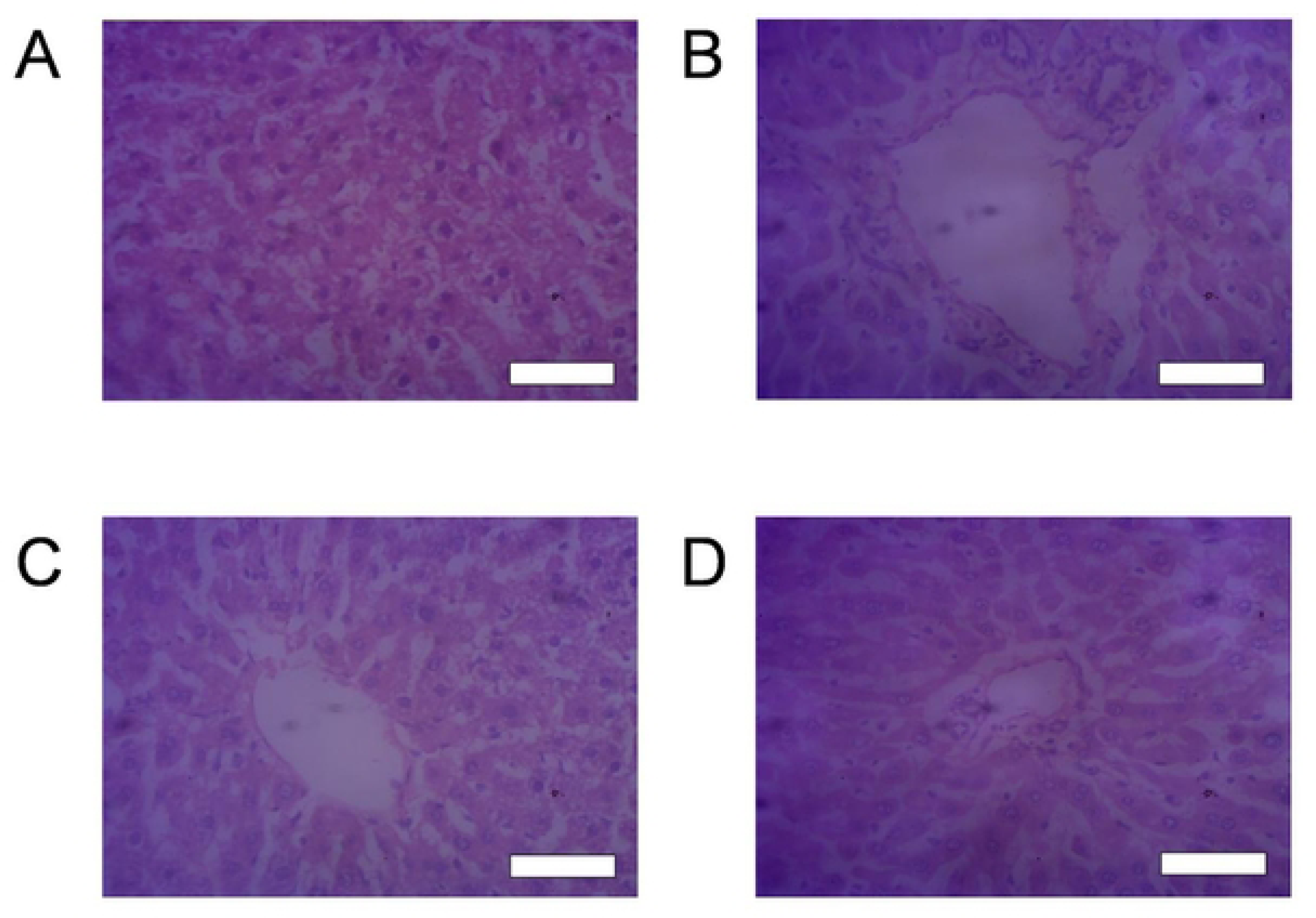
Histopathological examination of the effect GLE samples on cisplatin-treated liver using hematoxylin and eosin staining. (A) Rats treated with saline, (B) cisplatin-treated rats, (C) cisplatin-treated rats with crystalline GLE, and (D) cisplatin-treated rats with SMEDDS-GLE. All the rat livers were sectioned at 5 μm using microtome, and stained with hematoxylin and eosin. The stained sections were studied under light microscope at a magnification of × 40. Each bar represents 100 µm.

### 2.7. Nephroprotective function of GLE samples

SMEDDS-GLE is anticipated to boost the protective effects against cisplatin-induced kidney damage in rats. To assess the potential improvement of the protective effects of GLE, histological alterations, as well as plasma creatinine and BUN levels, which serve as indicators of nephrotoxicity, were examined in a rat model of acute nephrotoxicity induced by cisplatin. Cisplatin-induced nephrotoxicity is multifaceted etiology encompasses oxidative stress, inflammatory response, and apoptosis. Cisplatin can penetrate renal epithelial cells via organic cation transporter 2 (OCT2) and copper transporter 1 (Ctr1), leading to mitochondrial DNA damage and elevated levels of reactive oxygen species (ROS). This ultimately leads to tubular apoptosis and necrosis ^43,52^. The kidney tissues were examined under a light microscope after 72 h of cisplatin exposure. Histopathological alterations were observed using H&E staining (**Fig. 9**). **Fig. 9A** demonstrates that the kidney tissues in the control group displayed typical glomeruli, clearly defined cellular structures, and consistent tubules bordered with a single layer of epithelial cells. The kidney tissues in the group treated with cisplatin exhibited evident degenerative alterations, including atrophy of the epithelial lining, dilation of the tubules, necrosis of the tubules, congested glomeruli, and vacuolization of the cytoplasm in the tubular epithelial cells (**Fig. 9B**). The histological analysis of kidney tissues following repeated administration of GLE showed no notable distinctions compared to the group treated with cisplatin (**Fig. 9C**). In contrast, the group that received SMEDDS-GLE treatment showed normal morphology, with a significantly reduced level of glomerular degeneration and injury to tubular epithelial cells (**Fig. 9D**). The investigation of parenchymal injury, which is strongly associated with glomerular filtration, was conducted by assessing plasma creatinine and BUN levels as indicators of nephrotoxicity. The group treated with cisplatin (vehicle) had a notable (*p* < 0.01) elevation in creatinine and BUN levels when compared to the control group, suggesting serious damage to the kidney tissues (**Fig. 8**). Administering SMEDDS-GLE (75 mg GLE kg^−1^ for 10 days) significantly reduced the elevated levels of plasma creatinine and BUN by approximately 53% and 64% respectively, compared to the control group. There was a significant difference in the levels of these biomarkers between the control group and the group treated with SMEDDS-GLE (*p* = 0.048 and 0.020 for creatinine and BUN, respectively). The results indicate that the group treated with SMEDDS-GLE showed reduced pathological abnormalities caused by cisplatin, suggesting that GLE has enhanced nephroprotective effects when combined with SMEDDS-GLE. This finding is consistent with the better pharmacokinetic behavior of SMEDDS-GLE. Thus, improving the oral absorption of GLE by SMEDDS-GLE could enhance its positive benefits in preventing a range of disorders caused by free radicals in the body.

**Fig. 8.**
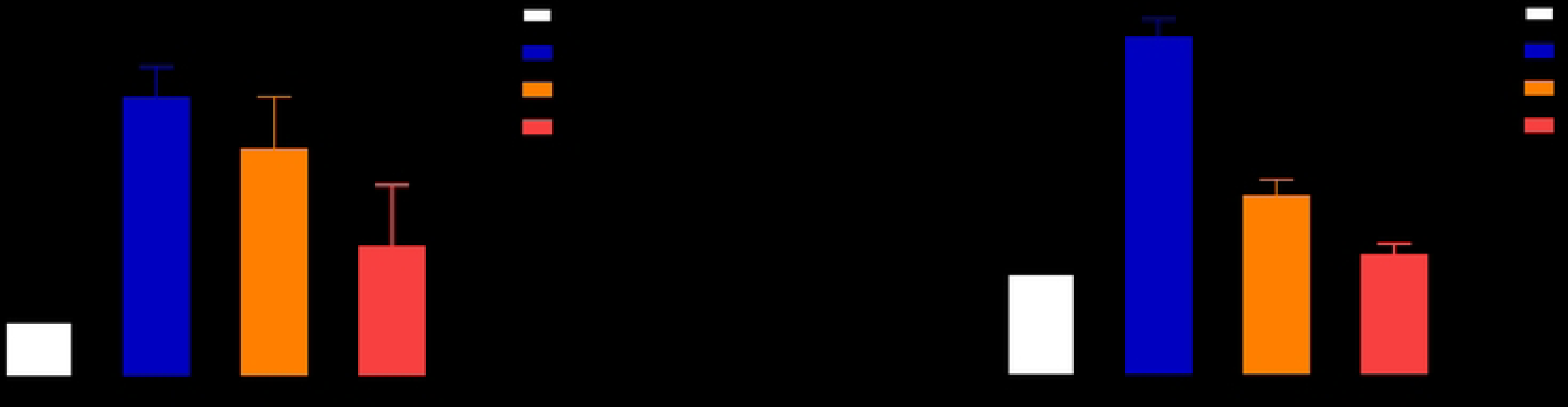
Nephrotoxic potential in a rat nephropathy model induced by cisplatin (7.5 mg/kg, *i.p.*). (A) Plasma creatinine and (B) BUN levels in rats with orally dosed GLE and SMEDDS-GLE (75 mg/kg, 10 days). **, *P*<0.01; ***, *P*<0.001 with respect to cisplatin-treated rats; ##, P< 0.01; ###, P<0.001, GLE vs. SMEDDS-GLE. Data represent mean ± S.E. of 6 experiments.

**Fig. 9.**
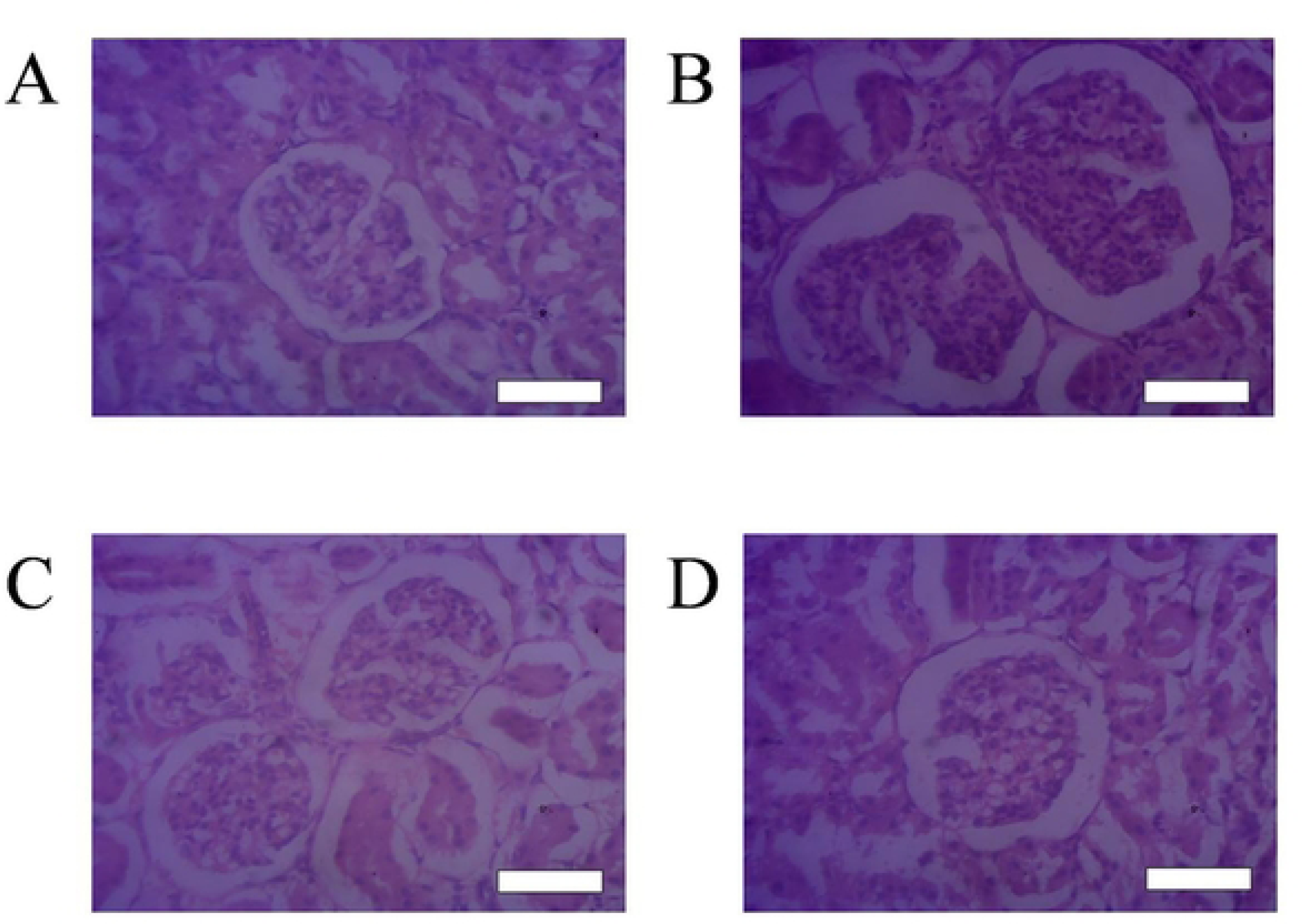
Histopathological examination of the effect GLE samples on cisplatin-treated kidney using hematoxylin and eosin staining. (A) Rats treated with saline, (B) cisplatin-treated rats, (C) cisplatin-treated rats with crystalline GLE, and (D) cisplatin-treated rats with SMEDDS-GLE. All the rat kidneys were sectioned at 5 μm using a microtome, and stained with hematoxylin and eosin. The stained sections were studied under a light microscope at a magnification of × 40. Each bar represents 100 µm.

## 3. Conclusion

The therapeutic applicability of GLE is limited because of its lipophilic composition, which also makes it appear to be poorly disseminated in GI fluid and to have very poor GI absorption of its clinically beneficial components. The current investigation aimed to create an enhanced self-nanoemulsifying drug delivery system (SMEDDS) formulation using a pseudo-ternary phase diagram and emulsion-forming capacity. The SMEDDS-GLE showed enhanced dispersion and disintegration of GLE in comparison to GLE. The combination of SMEDDS-GLE significantly improved the hepatoprotective properties of GLE in a rat model of acute liver damage. Furthermore, SMEDDS-GLE demonstrated significant nephroprotective properties of GLE in a rat model of acute kidney damage. The results suggest that strategically incorporating GLE into SMEDDS can enhance the physical and chemical characteristics and the nutritional advantages of GLE and other bioactive extracts.

## Abbreviations

ALP: alkaline phosphatase
ALT: alanine aminotransferase
ANOVA: analysis of variance
AST: aspartate aminotransferase
BUN: blood urea nitrogen
CKD: chronic kidney disease
DLS: Dynamic Light Scattering
FT-IR: fourier transform infrared spectroscopy
GLE: *Gynura procumbens* leaves extract
H&E: hematoxylin and eosin
HPLC: high performance liquid chromatography
MCT: medium chain triglyceride
ROS: reactive oxygen species
SMEDDS: self-emulsifying drug delivery system
TEM: transmission electron microscopy
UV: ultraviolet
ZP: zeta potential

## Declarations

### Conflict of interests

The authors declare no conflict of interest.

### Funding

This research received a partial grant from North South University, Dhaka, Bangladesh. (Grant Number: CTRG-21-SHLS-02).

### Author’s contributions

Manik Chandra Shill: Methodology, *in vivo* experiments, Validation, Formal analysis, Writing Review & Editing, resources; Md. Faisal Bin Jalal: Methodology, Investigation, *in vivo* experiments; Madhabi Lata Shuma: Methodology, Formal analysis, Data Curation, Writing - Review & Editing; Patricia Prova Mollick: Methodology, Investigation, *in vivo* experiments; Md. Abdul Muhit: Methodology, Data Curation, Writing - Review & Editing; Shimul Halder: Conceptualization, Project administration, Methodology, Validation, Formal analysis, Writing Original Draft, Review & Editing, Visualization.

## Acknowledgements

The authors wish to thank the authority of the Semiconductor Technology Research Center, the University of Dhaka, for their kindness in permitting the particle size analysis of the samples.

## Notes

### Competing Interest Statement

The authors have declared no competing interest.

## References

1. Lee, C. H. on. Prevalence of renal dysfunction in patients with cirrhosis according to ADQI-IAC working party proposal. Clin. Mol. Hepatol. 20, 185–191 (2014).

2. Shafiezadeh, R., Alavian, S. M., Namdar, H., Gholami-Fesharaki, M. & Esmaeili, S. S. Evaluating the efficacy of carum copticum seeds on the treatment of patients with nonalcoholic fatty liver disease: A multi-center, randomized, triple–blind, placebo-controlled clinical trial study. Hepat. Mon. 20, 1–8 (2020).

3. Chen, C. Y. et al. The prevalence and association of chronic kidney disease and diabetes in liver cirrhosis using different estimated glomerular filtration rate equation. Oncotarget 9, 2236–2248 (2018).

4. Bennadi, D. Self-medication: A current challenge. J. Basic Clin. Pharm. 5, 19 (2014).

5. Wang, T. Y. et al. Membrane oxidation enables the cytosolic entry of polyarginine cell-penetrating peptides. J. Biol. Chem. 291, 7902–7914 (2016).

6. Jomova, K. et al. Reactive oxygen species, toxicity, oxidative stress, and antioxidants: chronic diseases and aging. Arch. Toxicol. 97, 2499–2574 (2023).

7. Lobo, V., Patil, A., Phatak, A. & Chandra, N. Free radicals, antioxidants and functional foods: Impact on human health. Pharmacogn. Rev. 4, 118–126 (2010).

8. Chaachouay, N. & Zidane, L. Plant-Derived Natural Products: A Source for Drug Discovery and Development. Drugs Drug Candidates 3, 184–207 (2024).

9. Nasri, H., Baradaran, A., Shirzad, H. & Rafieian-Kopaei, M. New concepts in nutraceuticals as alternative for pharmaceuticals. Int. J. Prev. Med. 5, 1487–1499 (2014).

10. Basist, P. et al. Potential nephroprotective phytochemicals: Mechanism and future prospects. J. Ethnopharmacol. 283, 114743 (2022).

11. Mushtaq, S., Abbasi, B. H., Uzair, B. & Abbasi, R. Natural products as reservoirs of novel therapeutic agents. EXCLI J. 17, 420–451 (2018).

12. Sharifi-Rad, M. et al. Lifestyle, Oxidative Stress, and Antioxidants: Back and Forth in the Pathophysiology of Chronic Diseases. Front. Physiol. 11, 694 (2020).

13. Shin, G. H., Kim, J. T. & Park, H. J. Recent developments in nanoformulations of lipophilic functional foods. Trends Food Sci. Technol. 46, 144–157 (2015).

14. Hasler, C. M. Functional Foods: Benefits, Concerns and Challenges—A Position Paper from the American Council on Science and Health. J. Nutr. 132, 3772–3781 (2002).

15. Jobaer, M. A. et al. Phytochemical and Biological Investigation of an Indigenous Plant of Bangladesh, Gynura procumbens (Lour.) Merr.: Drug Discovery from Nature. Molecules 28, (2023).

16. Wardana, A. P. et al. Gynura procumbens NANOENCAPSULATION: A NOVEL PROMISING APPROACH TO COMBAT DENGUE INFECTION. Rasayan J. Chem. 16, 802–810 (2023).

17. Tan, J. N. et al. Antioxidant and Anti-Inflammatory Effects of Genus Gynura: A Systematic Review. Front. Pharmacol. 11, (2020).

18. Kaewseejan, N., Sutthikhum, V. & Siriamornpun, S. Potential of Gynura procumbens leaves as source of flavonoid-enriched fractions with enhanced antioxidant capacity. J. Funct. Foods 12, 120–128 (2015).

19. Ismail, M. A. H., Bahari, E. A., Ibrahim, F. S., Dasiman, R. & Amom, Z. Effects of gynura procumbens extract on liver function test of hypercholesterolemia induced rabbits. J. Teknol. 78, 49–54 (2016).

20. Vasconcelos, A. G. et al. Promising self-emulsifying drug delivery system loaded with lycopene from red guava (Psidium guajava L.): in vivo toxicity, biodistribution and cytotoxicity on DU-145 prostate cancer cells. Cancer Nanotechnol. 12, 1–29 (2021).

21. Nadzir, M. M., Idris, F. N. & Hat, K. Green synthesis of silver nanoparticle using Gynura procumbens aqueous extracts. AIP Conf. Proc. 2124, (2019).

22. Patel, P., Garala, K., Singh, S., Prajapati, B. G. & Chittasupho, C. Lipid-Based Nanoparticles in Delivering Bioactive Compounds for Improving Therapeutic Efficacy. Pharmaceuticals (Basel). 17, (2024).

23. Teja, P. K., Mithiya, J., Kate, A. S., Bairwa, K. & Chauthe, S. K. Herbal nanomedicines: Recent advancements, challenges, opportunities and regulatory overview. Phytomedicine 96, 153890 (2022).

24. Ki, B., Soo, J., Kang, S., Young, S. & Hong, S. Development of self-microemulsifying drug delivery systems (SMEDDS) for oral bioavailability enhancement of simvastatin in beagle dogs. 274, 65–73 (2004).

25. Mohsin, K. & Pouton, C. W. The influence of the ratio of lipid to surfactant and the presence of cosolvent on phase behaviour during aqueous dilution of lipid-based drug delivery systems. J. Drug Deliv. Sci. Technol. 22, 531–540 (2012).

26. Amresh, G., Agarwal, V. K. & Rao, C. V. Self microemulsifying formulation of Lagerstroemia speciosa against chemically induced hepatotoxicity. J. Tradit. Complement. Med. 8, 164–169 (2018).

27. Halder, S., Ogino, M., Seto, Y., Sato, H. & Onoue, S. Improved biopharmaceutical properties of carvedilol employing α-tocopheryl polyethylene glycol 1,000 succinate-based self-emulsifying drug delivery systems. Drug Dev. Ind. Pharm. 0, 1–32 (2018).

28. Chandra Shill, M., et al. Longevity Spinach (Gynura procumbens) Ameliorated Oxidative Stress and Inflammatory Mediators in Cisplatin-Induced Organ Dysfunction in Rats: Comprehensive in vivo and in silico Studies. Chem. Biodivers. e202301719 (2024) doi:10.1002/cbdv.202301719.

29. Mohammed, S. J. et al. Structural Characterization, Antimicrobial Activity, and in Vitro Cytotoxicity Effect of Black Seed Oil. Evidence-based Complement. Altern. Med. 2019, (2019).

30. Singh, V. K. & Seed, T. M. How necessary are animal models for modern drug discovery? Expert Opin. Drug Discov. 16, 1391–1397 (2021).

31. Bryda, E. C. The Mighty Mouse: the impact of rodents on advances in biomedical research. Mo. Med. 110, 207–211 (2013).

32. Aboraya, D. M., El Baz, A., Risha, E. F. & Abdelhamid, F. M. Hesperidin ameliorates cisplatin induced hepatotoxicity and attenuates oxidative damage, cell apoptosis, and inflammation in rats. Saudi J. Biol. Sci. 29, 3157–3166 (2022).

33. Perše, M. & Veceric-Haler, Z. Cisplatin-Induced Rodent Model of Kidney Injury : *Biomed Res*. Int. 2018, 1–29 (2018).

34. The Institutional Animal Care and Use Committee. Guidelines for Establishing Humane Endpoints in Animal Study Proposals. 1–4 (2019).

35. Quaresma, A. B., d’Acampora, A. J., Tramonte, R., Farias, D. C. de & Joly, F. S. Histological study of the liver and biochemistry of the blood of Wistar rats following ligature of right hepatic duct. Acta Cir. Bras. 22, 68–78 (2007).

36. Halder, S., Islam, A., Muhit, M. A., Shill, M. C. & Haider, S. S. Self-emulsifying drug delivery system of black seed oil with improved hypotriglyceridemic effect and enhanced hepatoprotective function. J. Funct. Foods 78, 104391 (2021).

37. Shah, A. V & Serajuddin, A. T. M. Development of Solid Self-Emulsifying Drug Delivery System (SEDDS) I: Use of Poloxamer 188 as Both Solidifying and Emulsifying Agent for Lipids. Pharm. Res. 29, 2817–2832 (2012).

38. Gurram, A. K. et al. Role of components in the formation of self-microemulsifying drug delivery systems. Indian J. Pharm. Sci. 77, 249–257 (2015).

39. Wu, J., Mei, P. & Lai, L. Microemulsion and interfacial properties of anionic/nonionic surfactant mixtures based on sulfonate surfactants: The influence of alcohol. J. Mol. Liq. 371, 120814 (2023).

40. Ed, M. & Julia, G. Toxicological Profile for Propylene Glycol. ATSDR’s Toxicol. Profiles (2002) doi:10.1201/9781420061888_ch136.

41. Feeney, O. M. et al. 50 years of oral lipid-based formulations: Provenance, progress and future perspectives. Adv. Drug Deliv. Rev. 101, 167–194 (2016).

42. Ghosh, A. et al. Stabilized Astaxanthin Nanoparticles Developed Using Flash Nanoprecipitation to Improve Oral Bioavailability and Hepatoprotective Effects. Pharmaceutics 15, (2023).

43. Ghosh, A. et al. Lysophosphatidylcholine-based liposome to improve oral absorption and nephroprotective effects of astaxanthin. J. Sci. Food Agric. 103, 2981–2988 (2023).

44. Boonsongrit, Y., Mueller, B. W. & Mitrevej, A. Characterization of drug-chitosan interaction by 1H NMR, FTIR and isothermal titration calorimetry. Eur. J. Pharm. Biopharm. 69, 388–395 (2008).

45. Pandey, M. M., Jaipal, A., Charde, S. Y., Goel, P. & Kumar, L. Dissolution enhancement of felodipine by amorphous nanodispersions using an amphiphilic polymer: insight into the role of drug–polymer interactions on drug dissolution. Pharm. Dev. Technol. 21, 463–474 (2016).

46. Mantawy, E. M., Tadros, M. G., Awad, A. S., Hassan, D. A. A. & El-Demerdash, E. Insights antifibrotic mechanism of methyl palmitate: Impact on nuclear factor kappa B and proinflammatory cytokines. Toxicol. Appl. Pharmacol. 258, 134–144 (2012).

47. Sookoian, S. & Pirola, C. J. Alanine and aspartate aminotransferase and glutamine-cycling pathway: their roles in pathogenesis of metabolic syndrome. World J. Gastroenterol. 18, 3775–3781 (2012).

48. Goeptar, a R., Scheerens, H. & Vermeulen, N. P. Oxygen and xenobiotic reductase activities of cytochrome P450. Crit. Rev. Toxicol. 25, 25–65 (1995).

49. Goltzman, D. & Miao, D. Alkaline Phosphatase. in Encyclopedia of Endocrine Diseases (ed. Martini, L.) 164–169 (Elsevier, 2004). 10.1016/B0-12-475570-4/00082-2.

50. El-Sheikh, S. M. A. et al. Gastroprotective, hepatoprotective, and nephroprotective effects of thymol against the adverse effects of acetylsalicylic acid in rats: Biochemical and histopathological studies. Saudi J. Biol. Sci. 29, 103289 (2022).

51. Onoue, S. et al. Physicochemical and biopharmaceutical characterization of amorphous solid dispersion of nobiletin, a citrus polymethoxylated flavone, with improved hepatoprotective effects. Eur. J. Pharm. Sci. 49, 453–460 (2013).

52. Yoshida, T., Kumagai, H., Kohsaka, T. & Ikegaya, N. Protective effects of relaxin against cisplatin-induced nephrotoxicity in rats. Nephron - Exp. Nephrol. 128, 9–20 (2014).

